# The risk of *Xylella fastidiosa* outbreaks will decrease in the Mediterranean olive-producing regions

**DOI:** 10.1101/2020.07.16.206474

**Authors:** M. Godefroid, M. Morente, T. Schartel, D. Cornara, A. Purcell, D. Gallego, A. Moreno, J.A. Pereira, A. Fereres

## Abstract

The bacterium *Xylella fastidiosa* (*Xf*) is a worldwide distributed invasive insect-borne plant pathogen, which causes lethal diseases to many economically-important crops including olives, citrus, almonds and grapes as well as numerous forest, ornamental, and uncultivated plants. The Mediterranean basin is the top supplier of olive oil with 93% of the world production and is consequently highly concerned about the recent invasion of *Xf* in Europe. Recently, bioeconomic models estimated putative losses induced by the spread of *Xf* across the European olive-producing area ranging from 1.9 to 5.2 billion euros over 50 years; however, such models did not take into account the insect vectors, which constitute a key driver of *Xf* spread. In the present study, we used bioclimatic species distribution models to predict the current and future climate suitability of the Mediterranean area for the main efficient or putative transmitters of *Xf* to olive (i.e. *Philaenus spumarius, Neophilaenus campestris* and *Aphrophora alni*). An important part of the total extent of the Mediterranean olive-producing area, mainly situated in southern Spain, Turkey and Greece, is predicted as currently poorly suitable for these vector species. Moreover, models forecast that nearly the totality of the Mediterranean olive-producing regions will likely become climatically little suitable for these vectors by 2050 due to climate change. In Europe, *Xf* outbreaks have occurred so far only in localities predicted as climatically suitable for these main vector species (e.g. the Apulia region of Italy) while the areas predicted as poorly suitable are still apparently *Xf*-free, which suggests that climate tolerances of vectors might play a main role in shaping *Xf* outbreaks patterns. This pattern highlights the crucial necessity of accounting for vectors when assessing risk of *Xf* outbreaks, and when considering vector-borne diseases in general. The risk maps presented here will have important practical application for the optimization of current and future strategies to control *Xf* in the Mediterranean region.

## Introduction

The bacterium *Xylella fastidiosa* Wells (1987) (*Xf*) is an insect-borne plant pathogen that induces severe diseases in a wide array of ornamental and agricultural crops including Pierce’s disease of grapevines (PD), olive quick decline syndrome (OQDS), almond leaf scorch (ALS) and citrus variegated chlorosis (CVC) among others (Delbianco et al., 2019). *Xylella fastidiosa* colonizes the xylem of plants and, when the bacterial load is sufficiently large reduces xylem sap circulation. This disruption of water transport along xylem vessels causes decline in yield and fruit quality and ultimately may lead to the death of plants (Hopkins, 1989).

Native to Americas, *Xf* has invaded many regions including Asia, the Middle East and Europe (EFSA PLH Panel., 2019). In 2013, an OQDS outbreak reported in the Apulia region of South-Eastern Italy represented the first confirmed introduction and establishment of *Xf* in Europe (Saponari et al., 2013). Thereafter, the bacterium has been detected on a diverse range of hosts in France, Spain, Portugal, Germany and Israel (https://gd.eppo.int/taxon/XYLEFA/distribution) (Fig 1). Although Xf has only recently been officially detected in Europe, epidemiological data indicate that the Xf introduction is likely an older event (Soubeyrand et al., 2018), suggesting that the pathogen could be widespread in southern Europe and that outbreaks might occur in new areas if favorable ecological conditions are met. Since no efficient treatment exists to cure infected trees, control strategies implemented in the European Union mainly rely on pathogen early detection, removal of susceptible hosts in localities experiencing outbreaks, and control of vectors and application of restrictive measures on planting material (EFSA PLH Panel., 2019).

**Figure 1.**
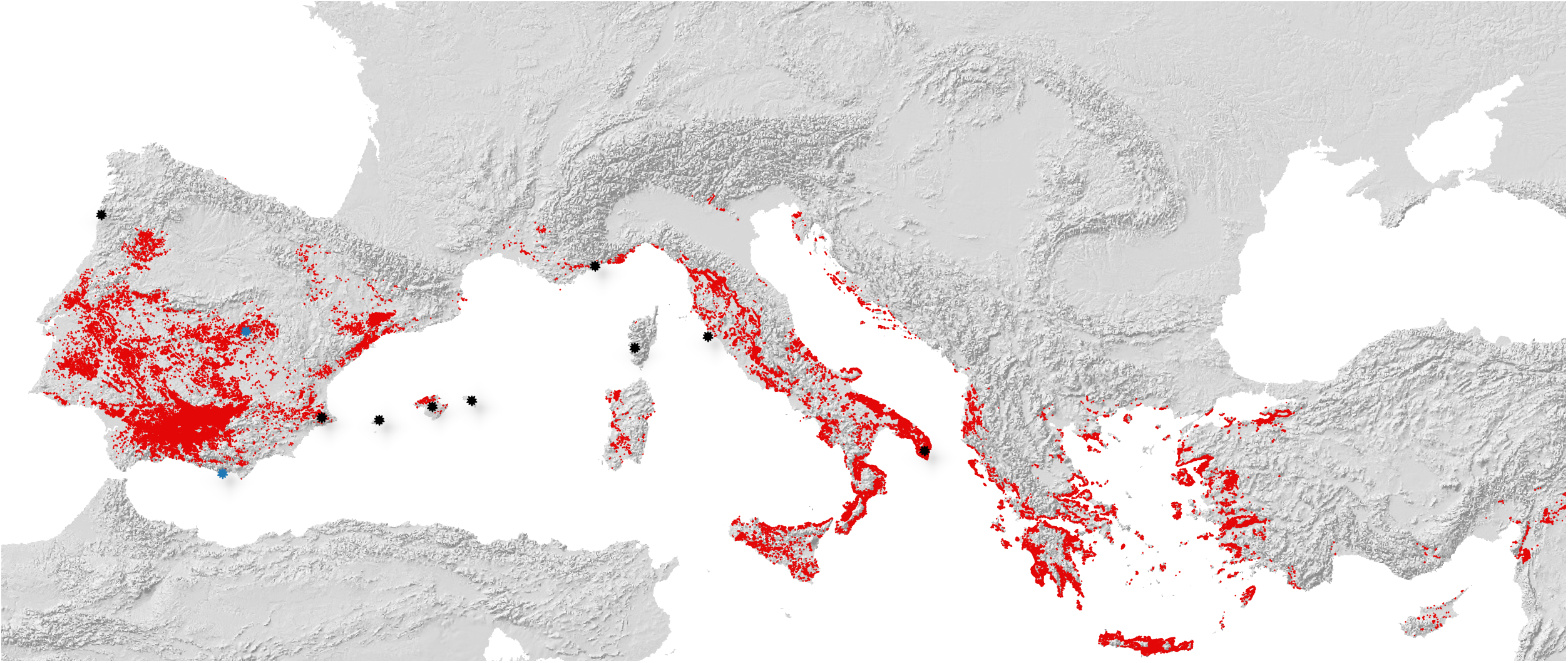
Distribution of olive groves and *Xylella fastidiosa* outbreaks in the Mediterranean area. The distribution of olive groves was extracted from the satellite-derived CORINE land cover database (CLC version 2018 - https://land.copernicus.eu/pan-european/corine-land-cover/clc2018). Minor and severe outbreaks of *Xylella fastidiosa* are represented by black and blue stars, respectively.

The worldwide economic impact of *Xf* is massive, e.g. annual losses caused by PD were estimated to reach ca. 104 million $US in California (Tumber et al., 2014) while the costs of removing trees affected by CVC were estimated at 120 million $US per year in Brazil (Bove & Ayres, 2007). Olive industry is severely threatened by *Xf* as recent OQDS outbreaks caused the death of hundreds of thousands of olive trees and induced an unprecedented economic crisis in the Apulian agriculture (Saponari et al., 2019). The Mediterranean basin is particularly concerned by *Xf* as it is the largest supplier of olive oil with ca. 93% of the world production (International Olive Council, www.internationaloliveoil.org/). In addition, the impact of *Xf* goes beyond purely economic losses, as olive cultivation is a centuries-old pillar of Mediterranean landscape, gastronomy and identity. Recently published bioeconomic models forecast that the olive-infecting Xf subspecies *pauca* might cause economic losses ranging from 1.9 to 5.2 billion euros over the next 50 years in Spain, Greece and Italy (Schneider et al., 2020); however, such models overlooked the pivotal factor mediating either Xf dispersal, or long-term establishment in a new area, i.e. insect vectors presence and abundance.

The *Xf* bacterium is transmitted to plants by insects feeding predominantly on xylem, such as sharpshooters (Hemiptera, Cicadellidae, Cicadellinae) and spittlebugs (Hemiptera, Aphrophoridae) (Cornara et al., 2019). Cicadas (Hemiptera, Cicadoidea) were also occasionally reported as vectors even though their involvement in *Xf* epidemiology, at least in Apulian olive orchards, is likely negligible (Cornara et al., 2020). The meadow spittlebug *Philaenus spumarius* L. (1978) (Hemiptera: Aphrophoridae) is currently the only epidemiologically relevant vector of *Xf* described in European outbreaks (Cavalieri et al., 2019; Cornara et al., 2017, 2020; Cruaud et al., 2018). Other two spittlebug species, i.e. *Philaenus italosignus* Drosopoulos & Remane (2000) and *Neophilaenus campestris* Fallén, (1805) (Hemiptera: Aphrophoridae) were found to be competent Xf vectors under experimental conditions (Cavalieri et al., 2019), in agreement with the “paradigm” that all the xylem-feeders are potentially vectors (Frazier, 1965). *Philaenus spumarius* mediates the late spring secondary transmission of *Xf* within olive orchards, as described by Cornara et al., (2017). On the other hand, given its sporadic presence on olive canopies, the contribution of *N. campestris* to OQDS epidemics is thought to be marginal or unlikely (Bodino et al., 2020; Cornara et al., 2017; Morente et al., 2018). Another species, *Aphrophora alni* (Fallén, 1805) (Hemiptera, Aphrophoridae), was found to be an important component of spittlebugs community on olive canopies in North-Western Italy (Liguria region) (Bodino et al., 2020). However, its vector competence has never been tested. Other xylem-feeders widely distributed in Europe - e.g. *Aphrophora* species, *Neophilaenus* species, *Cicadella viridis* L. (1758), *Lepyronia coleoptrata* L. (1758) or *Graphocephala fennahi* Young (1977) among others – are unlikely to play a role in Xf transmission to olive given their preference for other plant species (Cornara et al., 2019).

Growth of Xf in the plant xylem strongly depends on temperature conditions, with cold temperatures substantially reducing bacterial load in infected grapevines (“cold curing” phenomena; Feil & Purcell, 2001). Previous bioclimatic models aimed at predicting the risk of *Xf* outbreaks in Europe concluded that the Mediterranean countries were at high risk (Godefroid et al., 2019; Schneider et al., 2020). However, these predictions focused solely on the climatic tolerances of the bacterium and incompletely represented Xf epidemiology by omitting the vector ecology and distributions. Accounting only for the environmental tolerances of the bacteria itself is likely not sufficient as *Xf* outbreaks incidence also depends on vector field abundances, host preferences, dispersal patterns and transmission efficiencies (Daugherty & Almeida, 2009; Purcell, 1981). Transmission of Xf is positively correlated to vector abundance and the time they spend on the host, so severe *Xf* outbreaks may result from vectors with abundant or dense populations that reside for long periods on a Xf-host plant, despite having a relatively poor propensity (i.e. defined by Irwin & Ruesink (1986) as the probability of a vector transmitting a pathogen under field conditions) (Purcell, 1980).

The present study investigates the climate tolerances of the main efficient or putative vectors of *Xf* to olive and predicts the current and future climate suitability of European olive-producing regions for these vectors. To achieve this goal, we used bioclimatic species distribution models (SDMs) that establish a statistical relationship discriminating presence and absence of a species as a function of environmental conditions (Peterson et al., 2011). These models may serve to predict the potential distribution of a species in new space and time and are thus widely used in invasion risk analyses and assessments of global change effects on biodiversity (Peterson et al., 2011).

## Material and Methods

### Presence data

We collected distribution information for three efficient or putative vectors of *Xf* distributed in Europe namely *P. spumarius, N. campestris* and A. *alni. We* selected these species because they are the main known or putative vectors of Xf encountered in olive canopies from southern Europe (Antonatos et al., 2019; Bodino et al., 2020; Cornara et al., 2017; Lopes et al., 2014; Morente et al., 2018). Other xylem-feeding species that are rare in southern Europe or do not usually feed on olive trees, were not accounted for. We collected presence records in scientific literature and biodiversity databases such as the United States Federal Resource for Biological Occurrence Data (BISON - https://bison.usgs.gov/#home) and the public database Global Biodiversity Information Facility (GBIF - https://www.gbif.org/). Erroneous, ambiguous and non-reliable records were discarded from this dataset. We also assembled data collected by different Spanish plant protection agencies and research institutions (i.e. “*Instituto de Ciencias Agrarias*” at CSIC based in Madrid, Spain; “*Servicio de Sanidad Vegetal de la Junta de Andalucia*” based in Sevilla and Jaén, Spain, *Sanidad Agrícola Econex S.L*. based in Murcia province, Spain).

### Bioclimatic predictors

We downloaded five bioclimatic descriptors from the Worldclim 2.0 database at a resolution of 2.5 arc-min, which represent average historical climatic conditions during the 1970-2000 period (hereafter referred to as current climate conditions; Fick & Hijmans, 2017).These descriptors are the mean temperature of warmest quarter of the year (bio10), mean temperature of coldest quarter of the year (bio11), annual precipitation (bio12), precipitation of the warmest quarter of the year (bio18) and precipitation of the coldest quarter of the year (bio19). In addition, we also downloaded two descriptors representing air moisture conditions i.e. the Thornthwaite aridity index and the climatic index of moisture, from the ENVIREM database (Title & Bemmels, 2018). We selected all these descriptors for further ecological niche modelling because they are key climatic conditions that are ecologically-relevant for spittlebug occurrence, distributions, and survival (Karban & Strauss, 2004; Weaver & King, 1954; Whittaker & Tribe, 1996).

We predicted the climate suitability of the Mediterranean area for vectors by 2050 using future climate predictions reported in the sixth Assessment Report (AR6) of the Intergovernmental Panel on Climate Change (IPCC). To account for uncertainty in available forecasts of global change trends, we used future climates simulations from three global circulation models (GCM) i.e. the Model for Interdisciplinary Research on Climate version 6 MIROC6 (Watanabe et al., 2011), the Canadian Earth System Model version 5 (CanESM5; Swart et al., 2019) and the CNRM-ESM2-1 (Séférian et al., 2019). These simulations of future climates were extracted from the WorldClim 2.0 database, at a resolution of 2.5 arc-minutes. We considered two shared socioeconomic pathways i.e. scenarios SSP2-4.5 and SSP3-7.0 that are driven by different assumptions on future socioeconomics and represent moderate and relatively high greenhouse gas emissions respectively (O’Neill et al., 2014). Future values of the Thornthwaite aridity index and the climatic index of moisture were simulated using the *envirem* R package (Title & Bemmels, 2018), Worldclim monthly rasters of minimum and maximum temperature and precipitation, and rasters of extraterrestrial solar radiation obtained from the CGIAR-CSI website (http://www.cgiar-csi.org).

### Reduction of sampling bias in presence dataset

Before fitting models, we reduced occurrence autocorrelation in a climatic domain independently for each species, which is a recommended practice to reduce the putative negative impact of sampling bias on model predictions (Varela et al., 2014). We removed aggregation in a bi-dimensional space whose axes are the first two axes of a principal component analysis conducted on seven bioclimatic descriptors extracted at each pixel throughout the northern hemisphere. We divided each axis of the bi-dimensional space into 200 bins and then allowed only one occurrence per pixel in this 200 × 200 bins grid. A summary and maps of the distribution information available for each species is available in Appendix S1. The environmental filtering of presence records was purposely stringent and, consequently, a substantial amount of records was discarded. We adopted this conservative approach to reduce as much as possible the risk of omission error (i.e. false negative), which should be preferably avoided when assessing pest risk. In addition, models fitted with few climatically-filtered records tend to outperform models calibrated with large numbers of biased records (Varela et al., 2014).

### Absences and pseudo-absences for P. spumarius

The nature of absence data is one of the most crucial aspects in SDM (Lobo et al., 2010). While it is widely accepted that true absences (i.e. localities unsuitable for long-term establishment of the species under study) are preferable (Peterson et al., 2011), bioclimatic models are usually fitted using “pseudo-absences” (i.e. absences generated by the modeler) when true absences are lacking.

The current worldwide geographic distribution of *P. spumarius* is very well documented so we confidently generated pseudo-absences in geographic regions where this species is presumably absent or at least rare. We generated a maximum of 10,000 pseudo-absences in each of the following regions: (1) the southern parts of the Central valley of California; (2) the southeastern region of United States; (3) the north-eastern regions of Canada; (4) the high-altitude regions of Scandinavia and Iceland (elevation > 1000 meters) and Alps (elevation > 3000 meters); (5) the extremely dry Great Basin region across Nevada, Idaho and Oregon states (USA); (6) the Sahara desert (Appendix S2). We reduced autocorrelation of these pseudoabsence records in a climatic domain by applying the same approach used for filtering presence records. After climatic filtering, the final set of pseudo-absences encompassed 749 records. Although we cannot rule out that some of these pseudo-absences localities might be climatically suitable for *P. spumarius, we* assume that the majority of these pseudo-absences represent localities where the insect is relatively rare or absent, according to current knowledge on its distribution and biology.

We also compiled two sets of absences of *P. spumarius* recorded by Spanish plant security agencies (“*Servicio de Sanidad Vegetal*”) in southern regions of Spain. The first set of absences was retrieved from four years (2015-2018) of thorough and opportunistic field sampling performed in south-western Spain. This sampling consisted of visiting a total of 566 localities in various habitats (i.e. mostly natural habitats or olive groves with natural cover vegetation suitable for *P. spumarius* establishment). The sampling was focused on *P. spumarius* distribution and aimed at visiting different kinds of habitats and climatic regions in the province of Sevilla, Spain. A sampling point consisted in searching nymphs in natural vegetation during *ca*. 15 minutes in a radius of *ca*. 200 meters. Sampling was usually performed by two or three samplers with high experience and expert knowledge on spittlebugs ecology. Samplers followed main roads by car and performed sampling in areas with environmental conditions the most suitable as possible (abundant cover vegetation, good moisture conditions, suitable landscape, etc.). More details on this sampling survey are provided by Durán (2018).

The second set of absences was retrieved from an opportunistic field sampling performed in olive-cropping region of southern Spain in 2018-2019. This sampling consisted of visiting a total of 90 olive groves minimally managed with natural cover vegetation. A sampling point consisted in searching spittlebugs nymphs or adults in natural vegetation in olive groves and their surroundings during *ca*. 20 minutes in a radius of *ca*. 100-200 meters. Sampling was usually performed by one or two samplers with high experience and expert knowledge on spittlebugs ecology. More information on this sampling are provided by Torres (2020).

We assembled these two absence datasets (Appendix S3). Field-collected absences situated in the same pixel as a presence record on a bioclimatic raster at 10 arc-min resolution were discarded from this dataset to minimize the risk of using false absences. We purposely used a coarser resolution raster to filter absences to, as much as possible, avoid including false absences in the dataset. This filtered set of absences encompassed 197 records (referred as to “Abs-Spain1”). When fitting the full models (see below), we also discarded absence records situated inside the climate envelope defined by the 2.5th and 97.5th percentiles of the climate conditions extracted at *P. spumarius* presences. This final filtered set of absences of *P. spumarius* in southern Spain encompassed 93 records (referred as to “Abs-Spain2”; Appendix S3).

### Spittlebugs abundance data

An abundance dataset was collected in the province of Murcia, Spain, during spring 2019. Spittlebugs adults were collected using a sweeping net in 20 agricultural fields (i.e. 4 olive groves, 13 almond groves and 3 vineyards), twice a month from April to August 2019 (i.e. a total of 8 sampling dates for each field; Appendix S4). As natural vegetation was seldom in the core of most of fields because of soil tillage, sampling was also performed in their close surroundings. At each sampling date, a hundred sweeps were performed in each of three components of landscape i.e. cover herbaceous vegetation, the canopies of cultivated plants and the canopies of surrounding trees and shrubs. We summed the total spittlebug adults collected in each field and its surroundings (referred to as “*Murcia-abundance*” dataset) and also transformed these data into a binary presence-absence dataset (referred to as “*Murcia-binary*” dataset).

### Ensemble forecasting approach

Many SDM algorithms exist and may yield variable forecasts, warranting the usefulness of combining outputs of different techniques in an ensemble forecasting framework and providing a consensus prediction (Araújo & New, 2007). We modelled the climatic niche of Xf vectors using seven different modeling approaches i.e. generalized linear models (GLM), boosted regression trees (BRT), generalized additive models (GAM), random forest (RF), multiple adaptive regression splines (MARS) surface range envelope (SRE) and artificial neural networks (ANN). In model calibration, we equaled the weighted sum of presences and the weighted sum of pseudo-absences (Barbet-Massin et al., 2012). The predictive power of models was evaluated using the area under the curve (AUC) of the receiving operator curve (ROC; Fielding & Bell, 1997) and the True Skill Statistic (TSS; Allouche et al., 2006). We fitted models using 80 percent of data whereas the remaining 20 percent of records were kept for model evaluation. To account for uncertainty induced by cross-validation methods, we performed five replicates of every model. For each repetition, we processed a consensus prediction, which is the average predicted climatic suitability across the different modeling methods. To reduce uncertainty in projections, we combined predictions from models with TSS > 0.75 when processing the consensus. We weighted individual model contributions to the consensus predictions by their TSS scores. We evaluated the predictive power of each consensus prediction using the same part of the data reserved for model evaluation. Analyses were performed using the “biomod2” (Thuiller et al., 2016) package in R (R core Team, 2013).

### Bioclimatic predictors selection

The selection of the environmental dataset used in SDM has a crucial impact on the accuracy of potential geographic range predictions (Pliscoff et al., 2014). It is widely recognized that using ecologically-relevant climate descriptors that have a direct impact on the study species, enhances model transferability to novel space and time (Peterson et al., 2011). Also, as ecological data are often autocorrelated, a traditional random data partitioning into calibration/evaluation datasets may artificially inflate accuracy metrics. It is thus usually preferable to estimate model transferability using an evaluation dataset that is spatially or temporally independent from the calibration domain (Randin et al., 2006). Before fitting definitive full models, we thus applied a preliminary intercontinental transferability evaluation approach to select the best-predictive environmental dataset used in further models (Randin et al., 2006). This approach consists of fitting models using data from a region and evaluating their predictive power using evaluation data available for another geographic area. This approach therefore identifies proximal descriptors that have a direct causality to the biology of the study species and avoids potential issues concerning differences in predictor correlation structures among different geographic areas or time periods (Jiménez-Valverde et al., 2009).

We applied this approach to *P. spumarius* distribution data because this framework is particularly suited for a widely distributed polyphagous invasive species such as *P. spumarius* whose geographic distribution is wide, highly well-documented, at equilibrium (i.e. the species has likely reached most of areas climatically suitable in the biogeographic regions where it occurs) and minimally constrained by biotic factors (e.g. plant host distribution). We fitted models using only occurrences and pseudo-absences of *P. spumarius* situated in Americas and calculated the average predictive accuracy of consensus predictions (AUC and TSS) on the remaining presence/pseudo-absences data available for the rest of the world (“*eval1*”) as well as on the presence/absence data (i.e. “Abs-Spain1” absence dataset) collected by Spanish plant protection agencies in southern Spain (“*eval2*”). When performing this bioclimatic descriptor selection, environmental filtering of presences and pseudo-absences datasets was performed independently in America and in the rest of the world using the above-mentioned method. To select the best-predictive climate dataset, we discarded climate datasets associated with models with TSS < 0.6 and AUC < 0.8 for *eval1*. Afterwards, we kept the climate dataset associated with the model with highest AUC and TSS for *eval2* metrics. Using this approach, we tested twentyeight climate datasets representing different combinations of few-relevant bioclimatic descriptors. Since correlation between environmental descriptors may have a negative impact on model performance, we ensured that the climate datasets did not encompass highly-correlated variables (Pearson’s correlation index < 0.8 considering climatic values extracted at presence and pseudoabsence localities). At purpose, we fitted models using few bioclimatic variables to avoid model overfitting as much as possible (Randin et al., 2006). We adopted a such conservative approach according to the purpose of the study i.e. minimizing omission error and transferring model to new time and space.

### Pseudo-absences and bioclimatic predictors selection for N. campestris and A. alni

The geographic distribution of *N. campestris* and *A. alni* is not as wide as that of *P. spumarius* and is, consequently, much less suited to apply the above-described bioclimatic predictors selection framework. Indeed, *N. campestris* only occurs in Europe and the Middle East while *A. alni* is only distributed in Eurasia and eastern North America. As *N. campestris* and *A. alni* are spittlebugs closely genetically related to *P. spumarius*, with highly similar ecological characteristics (i.e. phenology, distribution patterns, etc.), we consider here that the physiology of these spittlebugs is likely constrained by similar bioclimatic descriptors. Then, the bioclimatic predictors dataset previously selected for *P. spumarius* was used to model the distribution of *N. campestris* and *A. alni*.

The geographic distribution of *N. campestris* and *A. alni* is not as well documented as that of *P. spumarius*, which precludes generating pseudo-absences based on expert-knowledge. For *N. campestris* and *A. alni, we* thus randomly generated 1,000 pseudo-absences situated outside the climate envelope defined by a surface rectangular envelope (SRE method with 0.025 as the quantile value; Barbet-Massin et al., 2012). To enhance model transferability, we generated pseudo-absences in wide geographic regions corresponding to where each species occurs (Appendix S5). For both species, we used the previously described method to remove removed autocorrelation of pseudo-absences in a climatic domain defined by the bioclimatic descriptors used for model calibration.

### Full model calibration/evaluation

We calibrated full models for the three species under study using the previously-selected set of bioclimatic predictors and applying the previously-described ensemble forecasting approach. In addition to the above-described model evaluation approach, we also calculated the AUC of the average of consensus predictions on the “*Murcia-binary*” dataset. We also fit a generalized linear model (Poisson family) to address the impact of the average predicted climatic suitability on the abundance of spittlebugs (“*Murcia-abundance*” dataset). These analyses were not performed for *A. alni* since this species was collected in just one of these 20 fields in Murcia province.

### Predictions in olive-cropping areas

We projected the average of consensus predictions in the Mediterranean area under current and future climate conditions. To assess the climate suitability of the European olive-cropping area for vectors, we downloaded the distribution of olive groves from the satellite-derived CORINE land cover database (CLC version 2018 - https://land.copernicus.eu/pan-european/corine-land-cover/clc2018) (Fig. 1). For all species, we implemented a threshold maximizing the sum of sensitivity and specificity of each consensus prediction to transform model forecasts into binary maps (Liu et al., 2013) and calculate the averaged percentage of olive-cropping localities predicted as climatically suitable.

## Results

### Bioclimatic descriptors selection

After applying the filters on both the *eval1* and *eval2* metrics, all models calibrated with climate variables representing temperature trends during the coldest month/quarter of the year (bio6/bio11) were discarded since they yielded very poor intercontinental transferability (Appendix S6). The selected best-predictive climate dataset encompassed the maximum temperature of the warmest quarter of the year (bio10) and the climatic index of moisture (Appendix S6). Ensemble models fitted with only American occurrences accurately predicted the distribution of *P. spumarius* in Europe (mean AUC ± standard deviation = 0.92 ± 0.01; mean TSS ± standard deviation = 0.77 ± 0.04) as well as the presence/absence dataset collected by Spanish plant protections agencies (mean AUC ± standard deviation = 0.8 ± 0.01; mean TSS ± standard deviation = 0.41 ± 0.13) (Appendix S6 & S7).

### Model predictions under current climate conditions P. spumarius

The consensus predictions yielded high predictive accuracy (mean AUC ± standard deviation = 0.98 ± 0.01; mean TSS ± standard deviation = 0.88 ± 0.03) indicating excellent model discrimination between presences and pseudo-absences. The average of consensus predictions predicted accurately the presence-absence data collected in Murcia province, South-Eastern Spain (AUC > 0.99). Similarly, the positive relationship between the average predicted climate suitability and abundance of *P. spumarius* adults in Murcia province, was highly significant according to a generalized linear model (*P-value* < 0.05).

The consensus prediction for *P. spumarius* indicated large parts of colder and humid olive-producing regions in southern France, Portugal, northern parts of Spain, Italy, Greece and Turkey are currently highly climatically suitable by (Fig. 2A). Noticeably, the Mediterranean areas where severe Xf outbreaks currently occur (i.e. Balearics islands, mid-altitude regions of Valencia province, north of Portugal, Corsica island, southeastern France, Apulia and Tuscany regions in Italy) yielded high climatic suitability for *P. spumarius* (Fig. 2A). Lower climate suitability was, however, predicted in olive-producing regions from southern Spain and coastal parts of Greece and Turkey (Fig. 2A). When applying the threshold maximizing the sum of sensitivity and specificity, 20.26 ± 1.65 % (mean ± standard deviation) of the total extent of the European olive-producing area was predicted as currently poorly suitable for *P. spumarius* (Table 1).

**Figure 2.**
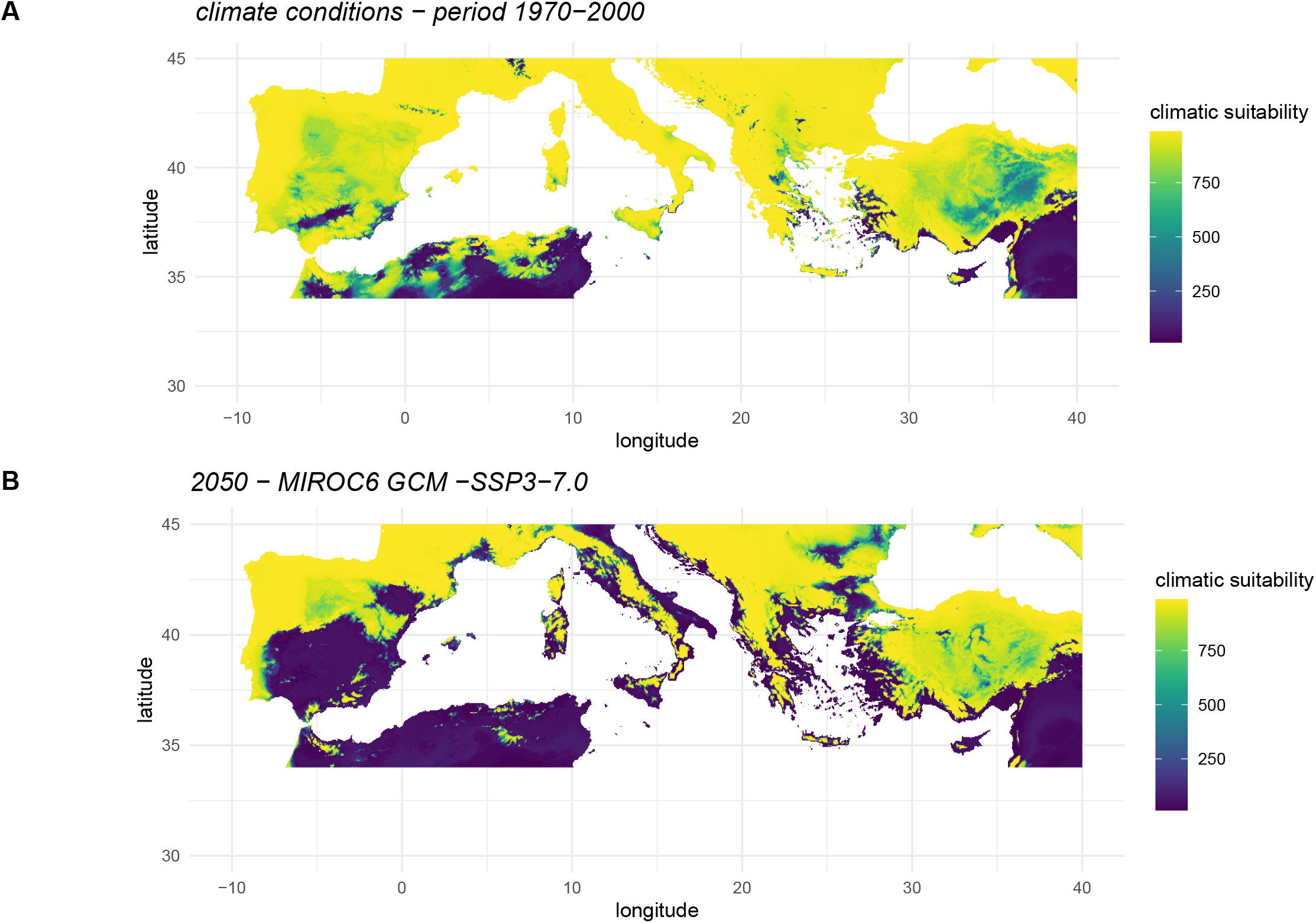
Consensus prediction of the potential distribution of *P. spumarius* under (A) current (1970-2000) and (B) future (2050) climate conditions. This consensus prediction was obtained using an ensemble forecasting method based on the modeling algorithms employed. The maps display the average predicted climate suitability by all models. We mapped predictions on future climate simulations (shared socioeconomic pathway SSP3-7.0) provided by the Model for Interdisciplinary Research on Climate version 6 MIROC6 available in the WorldClim database at a resolution of 2.5 arc-minutes.

**Table 1.**
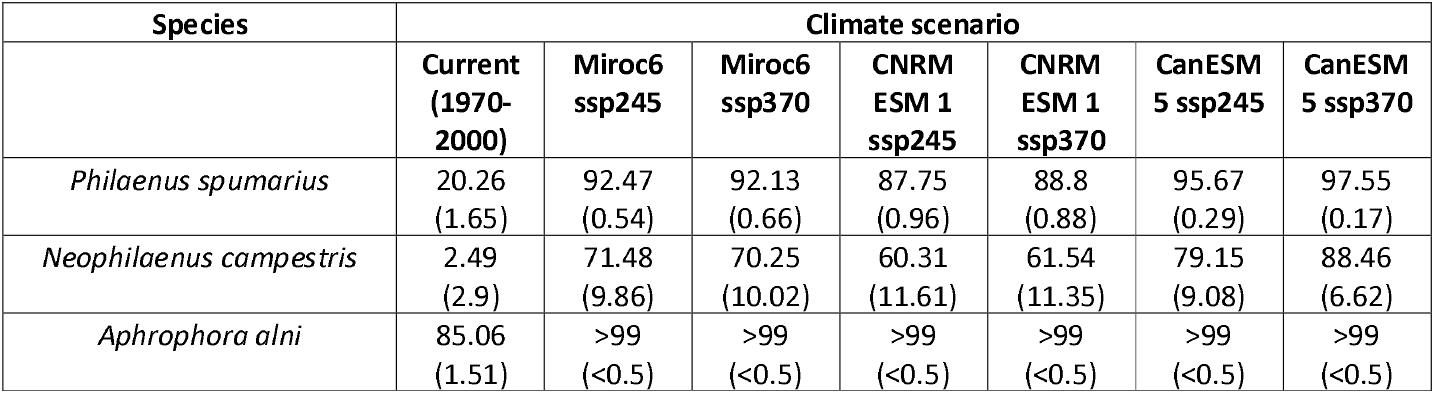
Mean and standard deviation (values in brackets) of the percentage (in %) of the total extent of European olive-producing regions predicted as poorly suitable by ensemble of bioclimatic species distribution models under current and future climate conditions (2050) for three efficient or putative vectors of *Xylella fastidiosa*.

### N. campestris and A. alni

The consensus predictions yielded high accuracy metrics for *A. alni* (mean TSS ± standard deviation = 0.96 ± 0.01; mean AUC ± standard deviation = 0.99 ± 0.004) and for *N. campestris* (mean TSS ± standard deviation = 0.92 ± 0.05; mean AUC ± standard deviation = 0.99 ± 0.01) indicating excellent models’ discrimination between presences and pseudoabsences. For *N. campestris*, the average consensus prediction predicted accurately the presence-absence data collected in Murcia province, South-Eastern Spain (AUC = 0.77).

Similarly, the positive relationship between the average predicted climate suitability and abundance of *N. campestris* in Murcia province was statistically significant according to a fitted generalized linear model (P-value < 0.05). Nearly the totality of European olive-producing regions was predicted as moderately suitable or poorly suitable for *A. alni* (Table 1; Fig. 3A). High climatic suitability for *A. alni* was predicted only in Mediterranean localities with relatively cold and humid climate conditions (Fig. 3A). Bioclimatic models predict large parts of Mediterranean basin – including most olive-producing regions - as moderately or highly climatically suitable for *N. campestris* (Table 1, Fig. 4A). Lower climatic suitability was predicted only in olive-producing regions from southern Spain and coastal parts of Greece and Turkey for this species (Fig. 4A). When applying the threshold maximizing the sum of sensitivity and specificity, 2.49 % and 85.06 % of the total extent of the European olive-producing area was predicted as currently poorly suitable for for *N. campestris* and *A. alni* respectively (Table 1).

**Figure 3.**
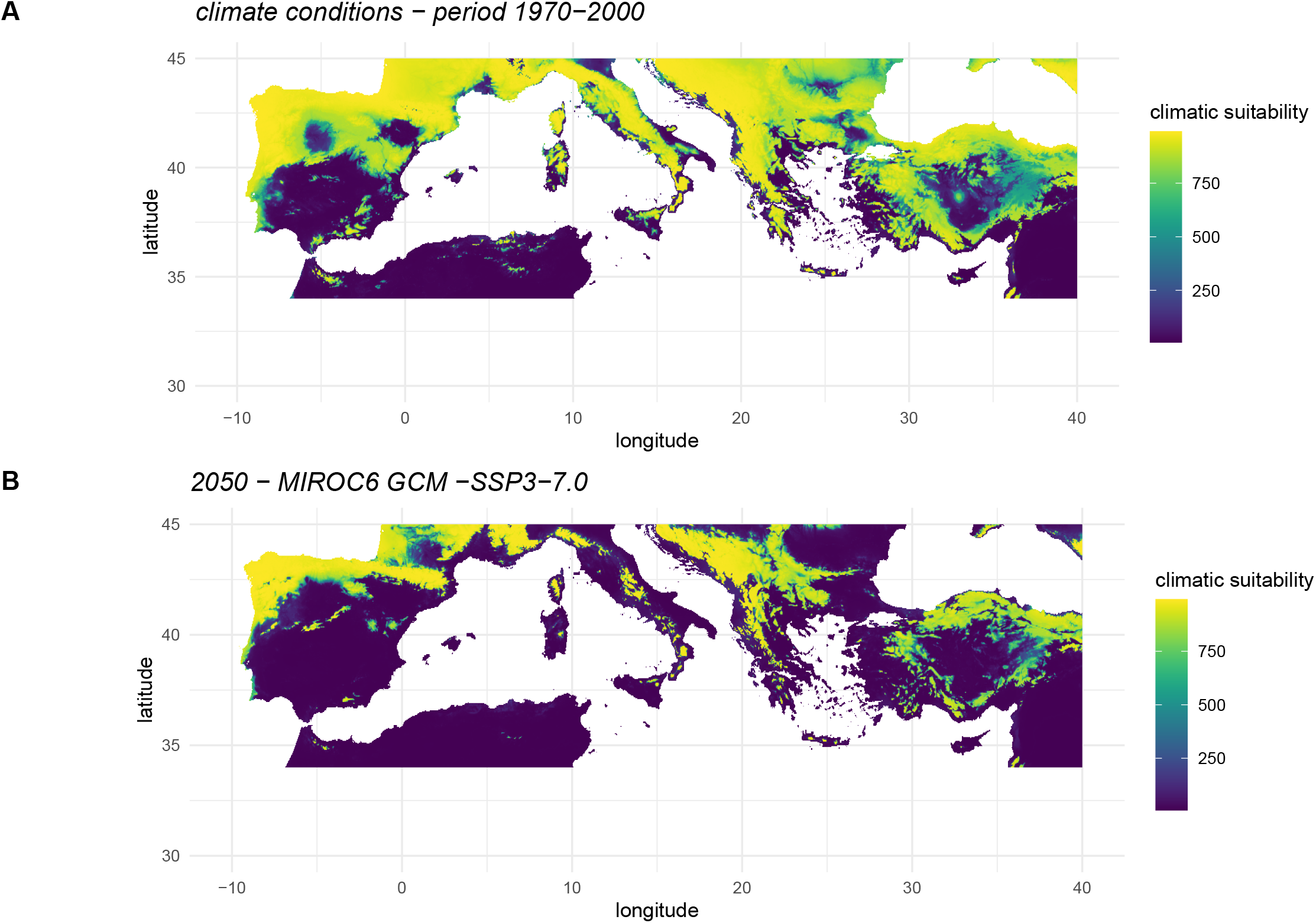
Consensus prediction of the potential distribution of *A. alni* under (A) current (1970-2000) and (B) future (2050) climate conditions. This consensus prediction was obtained using an ensemble forecasting method based on the modeling algorithms employed. The maps display the average predicted climate suitability by all models. We mapped predictions on future climate simulations (shared socioeconomic pathway SSP3-7.0) provided by the Model for Interdisciplinary Research on Climate version 6 MIROC6 available in the WorldClim database at a resolution of 2.5 arc-minutes.

**Figure 4.**
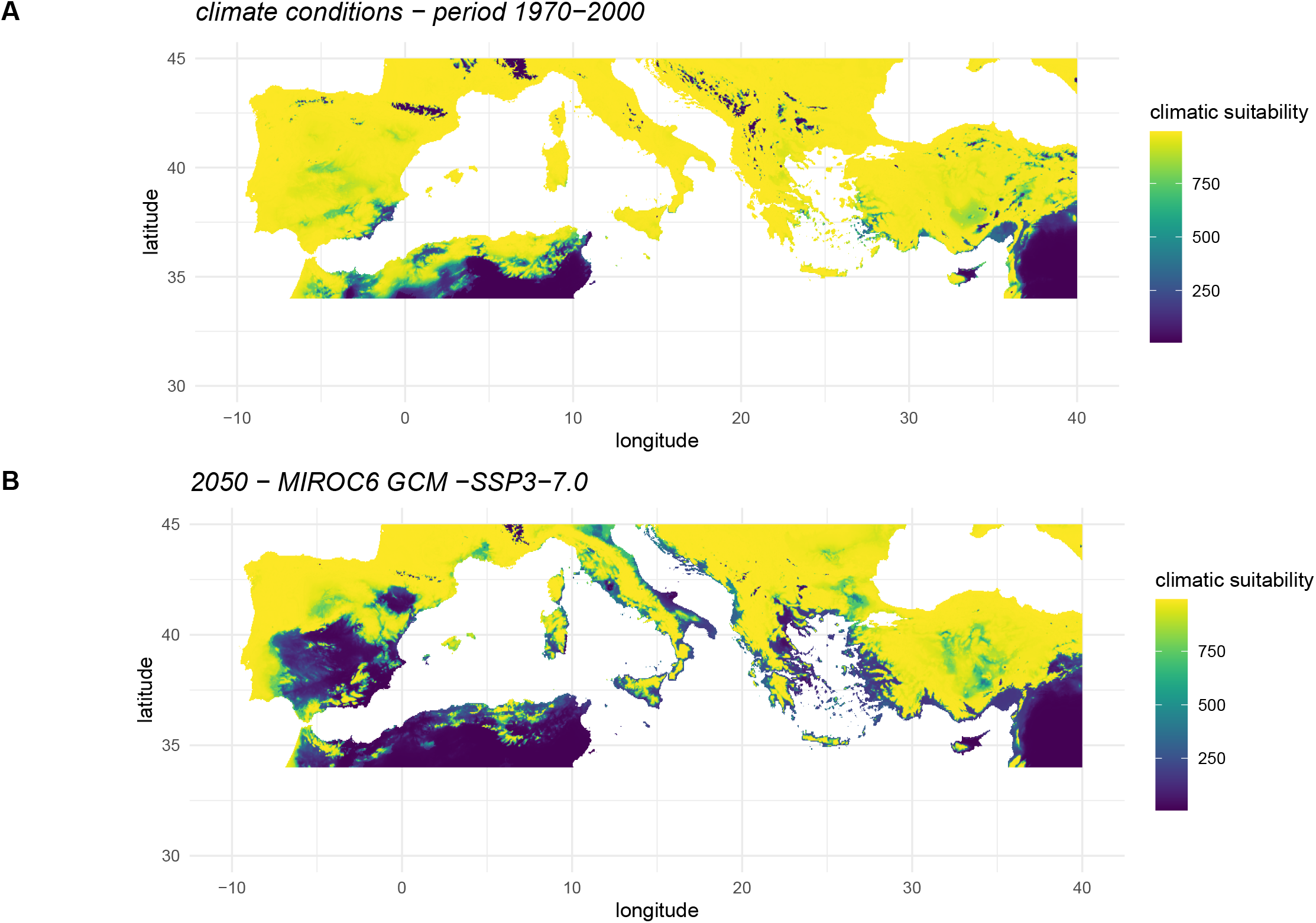
Consensus prediction of the potential distribution of *N. campestris* under (A) current (1970-2000) and (B) future (2050) climate conditions. This consensus prediction was obtained using an ensemble forecasting method based on the modeling algorithms employed. The maps display the average predicted climate suitability by all models. We mapped predictions on future climate simulations (shared socioeconomic pathway SSP3-7.0) provided by the Model for Interdisciplinary Research on Climate version 6 MIROC6 available in the WorldClim database at a resolution of 2.5 arc-minutes.

### Climate suitability under future climate conditions

Bioclimatic models predict that the Mediterranean olive-producing regions will experience a decrease in climatic suitability for all Xf vectors under study by 2050 (Table 1). Considering different future greenhouse gas emission scenarios, an minimum of ca. 87.75 %, 60.31 % and 99 % of the total extent of the European olive-producing area, are predicted to become poorly suitable for *P. spumarius, N. campestris* and *A. alni* by 2050, respectively (Table 1). Geographic areas that would become poorly suitable for these vectors by 2050 encompass most of lowlands in Greece, southern Portugal, southern and Central Spain and Italy including the Apulia region where severe Xf outbreaks currently occur (Figs. 2B, 3B, 4B).

## Discussion

### Climate suitability of the Mediterranean olive-producing regions for *Xylella fastidiosa* vectors

Several studies have addressed the environmental tolerances of Xf with the aim to predict which regions are at high risk in the Mediterranean basin (Godefroid et al., 2019; EFSA PLH Panel, 2019; Schneider et al., 2020). Considering mainly the mild winter conditions as favorable for bacterium systemic colonization and chronic infection establishment, bioclimatic models have predicted that the ongoing climate change will likely promote the emergence of *Xf* in most of the Mediterranean olive-producing area with dramatic socio-economic consequences (Schneider et al., 2020). However, such models overlooked the pivotal factor mediating *Xf* spread i.e. the distribution and abundance of insect vectors (Daugherty & Almeida, 2009; Purcell, 1981), and consequently tend to overestimate the risk in regions where efficient vectors are rare or absent.

The present study emphasizes the importance of accounting for vectors’ ecological characteristics when assessing *Xf* outbreak risk and highlights the risk of gathering unreliable predictions by excluding vectors when modelling the spread of a vector-borne plant pathogen. The availability of accurate forecasts of the distribution of pests is a prerequisite for the implementation of monitoring and control strategies in a rapidly changing world (Venette et al., 2010). Many harmful pests are currently experiencing rapid distribution shifts due to ongoing climate change or human activities (Parmesan, 2006) and the plant pathogen *Xf* is not an exception to this rule.

*Philaenus spumarius* is the only epidemiologically relevant Xf vector described in Europe (Cavalieri et al., 2019; Cornara et al., 2017, 2020; Cruaud et al., 2018) while the relative contribution of other species in *Xf* epidemiology is still debated and unclear. Although it is an ubiquitous and widespread species (Cornara et al., 2018), the models predict a marked spatial variation in climatic suitability throughout the European continent for *P. spumarius* i.e. models predict higher suitability in the colder regions of the Mediterranean basin while regions characterized by low humidity, limited annual precipitation and warm summer temperatures (e.g. lowlands in central and southern Spain, Greece and Turkey) display lower climatic suitability (Fig. 2A). Both other spittlebugs *A. alni* and *N. campestris* occur on olive canopies in Mediterranean regions and consequently should ideally be accounted for when designing monitoring plans and Xf control strategies. On the one hand, low climatic suitability was predicted for A. *alni* in most of the olive-producing regions of the Mediterranean basin (Table 1; Fig. 3A). This forecast is supported by field observations i.e. this spittlebug species is usually rare in most southern Europe olive groves (Antonatos et al., 2019; Morente et al., 2018) and is more abundant in colder and humid localities (Bodino et al., 2020). On the other hand, high climate suitability was predicted for *N. campestris* in most of the Mediterranean olive-cropping regions except in extremely warm regions of southern Spain (Table 1; Fig. 4A). This prediction is not surprising as *N. campestris* is the most common species found in most Mediterranean agrosystems together with *P. spumarius* (Antonatos et al., 2019; Bodino et al.,2020; Cornara et al.,2017; Morente et al., 2018).

Bioclimatic models suggest that climate change will likely be detrimental for the distribution of these three *Xf* vectors in the southern Europe. According to our models, the majority of European olive-producing regions – including areas where severe *Xf* outbreaks currently occur (e.g. Balearic Islands and Italy) - could become poorly suitable by 2050 for *P. spumarius, N. campestris* and *A. alni* (Figs. 2B, 3B, 4B; Table 1). These forecasts will have important practical applications when reassessing both the economic impact of *Xf* to the olive sector and the implementation of future control strategies. Indeed, bioeconomic models estimated massive putative economic losses due to spread of *Xf* in the European olive-producing regions over 50 years (Schneider et al., 2020); however, the present study mitigates these conclusions and calls for a reassessment of future Xf impact in the Mediterranean olive-cropping countries.

### Vectors-bacterium-climate interactions

According to Purcell (1981), the frequency and intensity of Xf outbreaks are positively correlated to abundance of efficient vectors in the field, the time they spend on hosts, their transmission efficiency and the bacterial load in infected plants. Interestingly, on the one hand, Xf outbreaks are currently occurring in European regions where high densities of putative vectors occur (Cornara et al., 2017; Cruaud et al., 2018; Lorente et al., 2018), and predicted as highly climatically suitable for *P. spumarius* and *N. campestris* (i.e. Balearic Islands, mid-altitude regions of Valencia province, north of Portugal, Corsica islands lowlands, southeastern France, central and southern Italy; Figs. 2 & 4). On the other hand, no severe Xf outbreak has ever occurred in southern areas of the Iberian Peninsula, Greece or Turkey (Fig. 1), even though these areas were predicted as suitable for the establishment of different Xf bacterium subspecies (Godefroid et al., 2019; Schneider et al., 2020). One hypothesis to explain this pattern is that climate conditions preclude the establishment of local abundant populations of efficient vectors in these regions and, consequently, drastically reduce probability of *Xf* transmission to plants and subsequent spread of the pathogen. While densities ofXfvectors are considered as relatively low in these warm and dry regions (Antonatos et al., 2019; Durán, 2018; Morente et al., 2018; Torres, 2020), we are aware that current data are not sufficient to prove undoubtedly the veracity of this hypothesis. To properly verify this hypothesis, we should ideally need to assess that Xf was introduced in these regions and did not spread because of the lack of abundant vector populations. Although it is highly likely that the bacterium *Xf* was long ago introduced in all regions of Europe (Soubeyrand et al., 2018) and that the incidence of outbreaks is directly associated with the local abundance of vector populations, further research should be needed to explore this hypothesis. Also, the impact of landscape structure, host plant distribution, agricultural practices and human disturbances should be accounted for in the modeling effort as all these factors interact to shape spittlebug densities in agricultural fields (Santoiemma et al., 2019).

### Reliability of models and perspectives

These models presented here were calibrated using an unprecedented distribution dataset based on an extensive mining of scientific literature and biodiversity databases as well as a thorough field-collected dataset from the southern Iberian Peninsula. Models were calibrated using few ecologically-relevant climate descriptors, which is a conservative approach to avoid overfitting, reduce omission risk and enhance model transferability (Randin et al., 2006). This conservative approach was preferred since our main goal was to assess the potential spread of a vector-borne plant pathogen with a devastating impact in economically-important agricultural regions.

The models identified summer temperature, annual rainfall and moisture as good predictors of the range of *P. spumarius* (Appendix S6), which is consistent with existing knowledge on the biology of the meadow spittlebug (Cornara et al., 2018; Karban & Strauss, 2004; Weaver & King, 1954). For instance, low humidity and high temperature were proven to be detrimental to all life stages of *P. spumarius* under laboratory conditions (Weaver & King, 1954). Summer temperature and humidity were repeatedly found to be good predictors of the abundance of the meadow spittlebug in North America (Karban & Strauss, 2004; Weaver & King, 1954) as well as other spittlebugs in Europe (Whittaker, 1971; Whittaker & Tribe, 1996). Similarly, the recent increase of summer temperature and drought conditions has been suggested to be the most likely cause of the currently observable decline of meadow spittlebug densities along the California coast (Karban & Huntzinger, 2018; Karban & Strauss, 2004).

Overall, the models yielded excellent accuracy metrics and the predictions agree with expert knowledge on the distribution pattern of *P. spumarius* in Spain i.e. this spittlebug is particularly rare in warm and dry lowlands of southern Spain, such as the Guadalquivir basin in Andalucía (Durán, 2018). Moreover, a highly significant statistical relationship was found between the number of *P. spumarius* adults collected in agricultural fields of South-Eastern Spain (Murcia) and model-predicted climatic suitability. We are aware that the scarcity of *P. spumarius* in agricultural regions of dry and warm lowlands of southern Spain might be explained by additional factors such as landscape structure, host plant distribution, agricultural practices and human disturbances. Olive cropping in southern Spain is particularly intensive (i.e. frequent soil tillage, weeds removal and application of chemical inputs), which likely negatively impacts populations of spittlebugs, similarly to what has been observed in Italian vineyards (Santoiemma et al., 2019). The present study suggests, however, that local climate conditions likely contribute in shaping this pattern as bioclimatic models fitted with only American occurrences predicted accurately field-collected data in southern Spain. The meadow spittlebug is a very widespread and highly polyphagous organism that occurs in nearly all habitats so that its absence or scarcity from large parts of South-Eastern United States and non-coastal regions of California is likely climate-driven (Karban & Strauss, 2004; Weaver & King, 1954). As a consequence, models fitted with available distribution data for *P. spumarius* in North America certainly give valuable information on its climatic preferences. Similarly, comparison of American and European distribution patterns of *A. alni* (i.e. this species occurs in cold and humid regions in both continents) strengthens the reliability of forecasts for this species. Unfortunately, model predictions for *N. campestris* are more uncertain as this species is only present in Europe and the Middle East so available distribution data are not as much informative as for the other spittlebugs under study and preclude a strict comparison of climate preferences among spatially-independent geographic areas. However, the absence of any *Neophilaenus* species in very warm regions of North America, the relative scarcity of *N. campestris* in southern Spain as well as the frequent cooccurrence of *N. campestris* and *P. spumarius* in Europe and the Middle East suggest that both species likely display similar climatic tolerances. Moreover, bioclimatic models predicted very accurately the abundance patterns of *N. campestris* in Murcia province (South-Eastern Spain), which strengthens the reliability of forecasts.

We urge caution when interpreting SDM outputs since such correlative approach only depicts the realized climatic niche of species i.e. a subset of their full environmental tolerances constrained by biotic interactions and dispersal constraints (Hutchinson, 1957). We also would like to stress that bioclimatic models do not account for micro-scale climatic factors that could increase suitability of some areas (e.g. crop irrigation, vicinity of a river, etc.). We suggest that calibration of mechanistic models based on physiological data is a key research path to provide better estimates of the potential climate suitability of olive-cropping regions for pests in a changing world (Ponti et al., 2014). Finally, we remind that the models fitted here are only based on bioclimatic predictors and thus just provide estimates of continental-scale climatic suitability. The distribution of spittlebugs is shaped by many additional factors that were not accounted for in the present study e.g. biotic interactions, dispersal range, plant-insect interactions, insects’ feeding behavior, cropping practices (e.g. soil tillage, use of phytosanitary products, planting of cover crops, soil nitrogen content) and landscape structure (e.g. the distribution of oversummering hosts). Also, importantly, the nature of herbaceous vegetation is crucial for nymphal development of spittlebugs - e.g. *P. spumarius* nymphs preferentially feed on Asteraceae and Fabaceae while nymphal instars of *N. campestris* preferentially develop on Poaceae (Bodino et al., 2020; Dongiovanni et al., 2019; Mazzoni, 2005; Morente et al., 2018) – and, then, should ideally be accounted for to improve the reliability of further risk assessments of Xf outbreaks.

### Conclusions

The present study provides estimates of the potential suitability of the Mediterranean basin under current and future climate conditions for the main vectors of Xf. Bioclimatic models highlight that some of the most economically-important olive-cropping regions situated in Spain, Greece and Turkey are not climatically optimal for the establishment of the main vectors of Xf. This finding might explain why Xf outbreaks have never occurred in these regions despite the bacterium *Xf* being likely widespread throughout Europe. Importantly, models predict that a substantial part of the Mediterranean olive-producing regions might become poorly suitable for the main Xf vectors by 2050 due to ongoing climate change. The maps provided here are easily interpretable tools that provide valuable information to design monitoring strategies and might enhance communication between the different actors involved in emerging pests control such as politics, decision-makers, funders, citizens and farmers.

## Supporting information

Appendi S6

## Acknowledgments

We are grateful to Servicio de Sanidad Vegetal de la Junta de Andalucia for providing data sets on the geographical distribution of spittlebugs in the Andalucia Region. We also thank Sanidad Agrícola Econex S.L for sharing data. This work has been financially supported by the Spanish Ministerio de Ciencia, Innovación y Universidades, Grant/Award Number: AGL2017-89604-R and the European Union Horizon 2020 research and innovation program under grant agreements no. 635646 POnTE (Pest Organisms Threatening Europe), and no. 727987 XF-ACTORS (Xylella Fastidiosa Active Containment Through a multidisciplinary-Oriented Research Strategy). The first author of this study was funded by the fellowship “Ayudas destinadas a la atracción de talento investigador de la Comunidad de Madrid”. Daniele Cornara participation in this work was supported by a research grant in the frame of European Union’s Horizon 2020 research and innovation programme under the Marie Skłodowska-Curie grant agreement No 835732 XYL-SPIT. We acknowledge the World Climate Research Programme, which, through its Working Group on Coupled Modelling, coordinated and promoted CMIP6. We thank the climate modeling groups for producing and making available their model output, the Earth System Grid Federation (ESGF) for archiving the data and providing access, and the multiple funding agencies who support CMIP6 and ESGF.

## Supplementary Information

**Appendix S1 (a-d)** Summary and mapping of the distribution information available for every species under study. All presence records available before environmental-filtering were represented by a black dot on maps.

**Appendix S2** Worldwide distribution of the pseudo-absences generated for *Philaenus spumarius*. Presences and pseudo-absences are colored in black and red respectively.

**Appendix S3** Mapping of the *P. spumarius* presence/absence records sampled in southern Spain by Spanish plant protection regional agencies. Presences and pseudo-absences are colored in black and red, respectively.

**Appendix S4** Map of presence-absence data in Murcia province, South-Eastern Spain. Black and red dots indicate presence and absence of vectors respectively.

**Appendix S5** Extent area where pseudo-absences were generated for each vector species under study

**Appendix S6** Model intercontinental accuracy metrics for *Philaenus spumarius*.

**Appendix S7** Consensus prediction of the potential distribution of *Philaenus spumarius* under current (1970-2000) climate conditions predicted by models were calibrated using only the occurrences and pseudo-absences available in North America. This consensus prediction was obtained using an ensemble forecasting method based on the modeling algorithms employed.

