## Appendi S6 for "The risk of *Xylella fastidiosa* outbreaks will decrease in the Mediterranean olive-producing regions"

| **Climate descriptor #** | **Climate descriptors** | **Eval1** | | **Eval2** | |
| --- | --- | --- | --- | --- | --- |
|  |  | AUC | TSS | AUC | TSS |
| 1 | bio10,bio18,bio19 | 0.82 | 0.45 | 0.74 | 0.07 |
| 2 | bio10,bio11,bio19 | 0.57 | -0.04 | 0.76 | 0.28 |
| 3 | bio11,bio18,bio19 | 0.36 | -0.12 | 0.58 | 0.12 |
| 4 | bio10,bio11,bio12 | 0.67 | -0.02 | 0.78 | 0.18 |
| 5 | bio10,bio11,bio18,bio19 | 0.67 | 0.03 | 0.74 | 0.17 |
| 6 | bio10,bio11,bio19 | 0.57 | 0.02 | 0.76 | 0.31 |
| 7 | bio10,bio11,bio18 | 0.75 | 0.16 | 0.79 | 0.05 |
| 8 | bio5,bio19 | 0.87 | 0.58 | 0.69 | 0.28 |
| 9 | bio5,bio18 | 0.88 | 0.54 | 0.57 | 0.15 |
| 10 | bio5,bio12 | 0.87 | 0.61 | 0.73 | 0.26 |
| 11 | bio6,bio19 | 0.30 | -0.41 | 0.81 | 0.002 |
| 12 | bio6,bio12 | 0.22 | -0.26 | 0.69 | 0.15 |
| 13 | bio6,bio18 | 0.19 | -0.25 | 0.35 | 0.02 |
| 14 | bio10,bio19 | 0.93 | 0.59 | 0.75 | 0.37 |
| 15 | bio10,bio18 | 0.73 | 0.52 | 0.48 | 0.05 |
| 16 | bio10,bio12 | 0.87 | 0.55 | 0.76 | 0.37 |
| 17 | bio11,bio19 | 0.33 | -0.24 | 0.77 | -0.001 |
| 18 | bio11,bio12 | 0.33 | -0.17 | 0.63 | -0.002 |
| 19 | bio11,bio18 | 0.3 | -0.06 | 0.42 | 0.002 |
| **20** | **bio10,moisture** | **0.92** | **0.78** | **0.8** | **0.41** |
| 21 | bio10,aridity | 0.94 | 0.69 | 0.63 | 0.05 |
| 22 | bio11,moisture | 0.4 | -0.12 | 0.61 | -0.01 |
| 23 | bio11,aridity | 0.34 | -0.04 | 0.49 | -0.06 |
| 24 | bio10,bio12,moisture | 0.88 | 0.5 | 0.79 | 0.43 |
| 25 | bio10,bio12,aridity | 0.87 | 0.56 | 0.8 | 0.2 |
| 26 | bio11,bio12,moisture | 0.49 | 0.04 | 0.72 | 0.23 |
| 27 | bio11,bio12,aridity | 0.37 | -0.12 | 0.77 | 0.24 |
| 28 | bio10,bio19,aridity | 0.85 | 0.43 | 0.78 | 0.2 |

**Appendix S5** Intercontinental evaluation metrics of ensemble forecast predictions. Models were fitted using presence records of *Philaenus spumarius* available in North America and were evaluated using two datasets : (1) all remaining occurrences and pseudo-asbences of *P. spumarius* distributed in the rest of the world (*eval1*) and (2) a presence-absence dataset field collected by Spanish Plant protection agencies in southern Spain (*eval2*). The best-predictive climate descriptor is bolded.
