## Supplementary figures and images for "The risk of *Xylella fastidiosa* outbreaks will decrease in the Mediterranean olive-producing regions"

### Appendix S1a

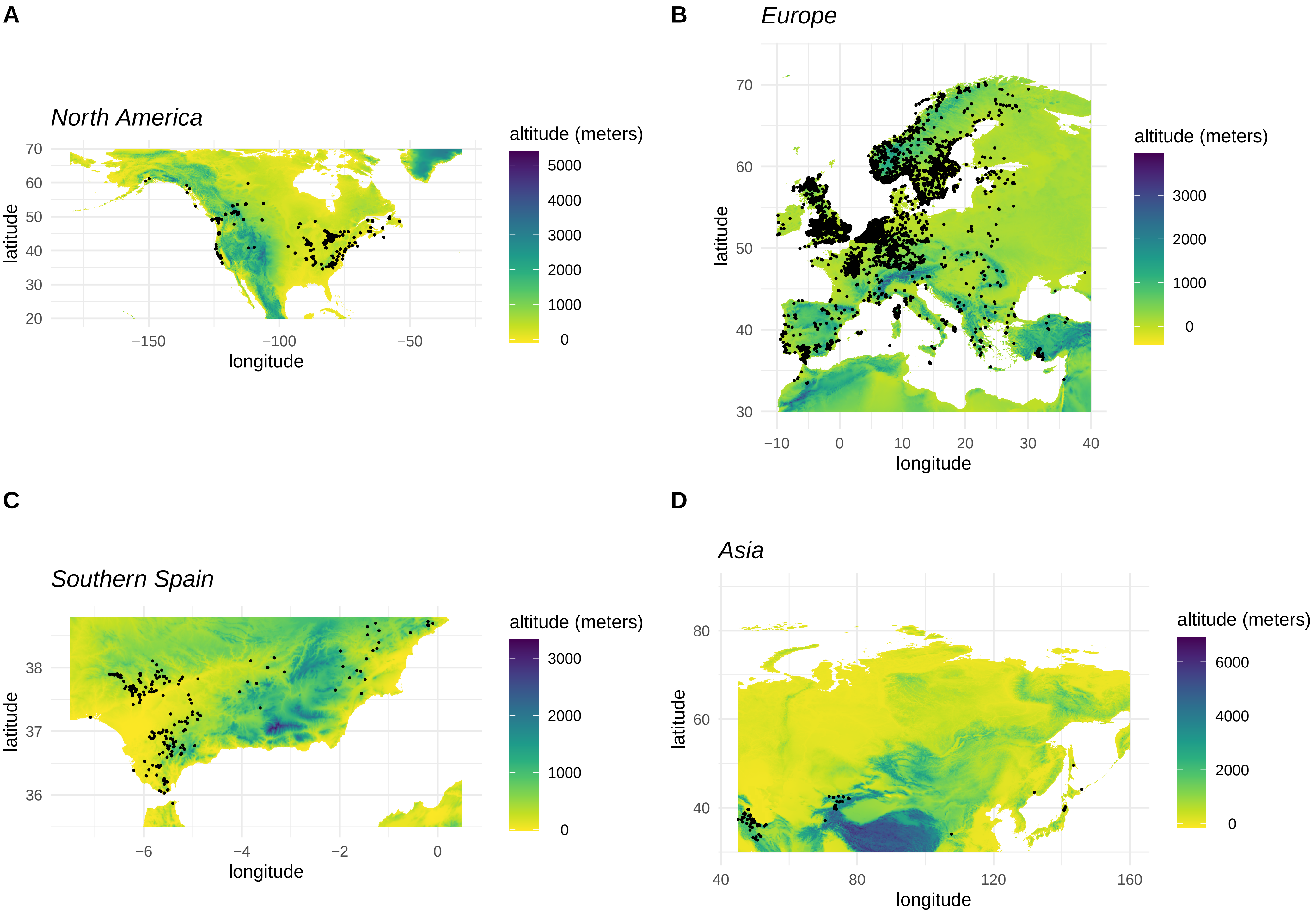

### Appendix S1b

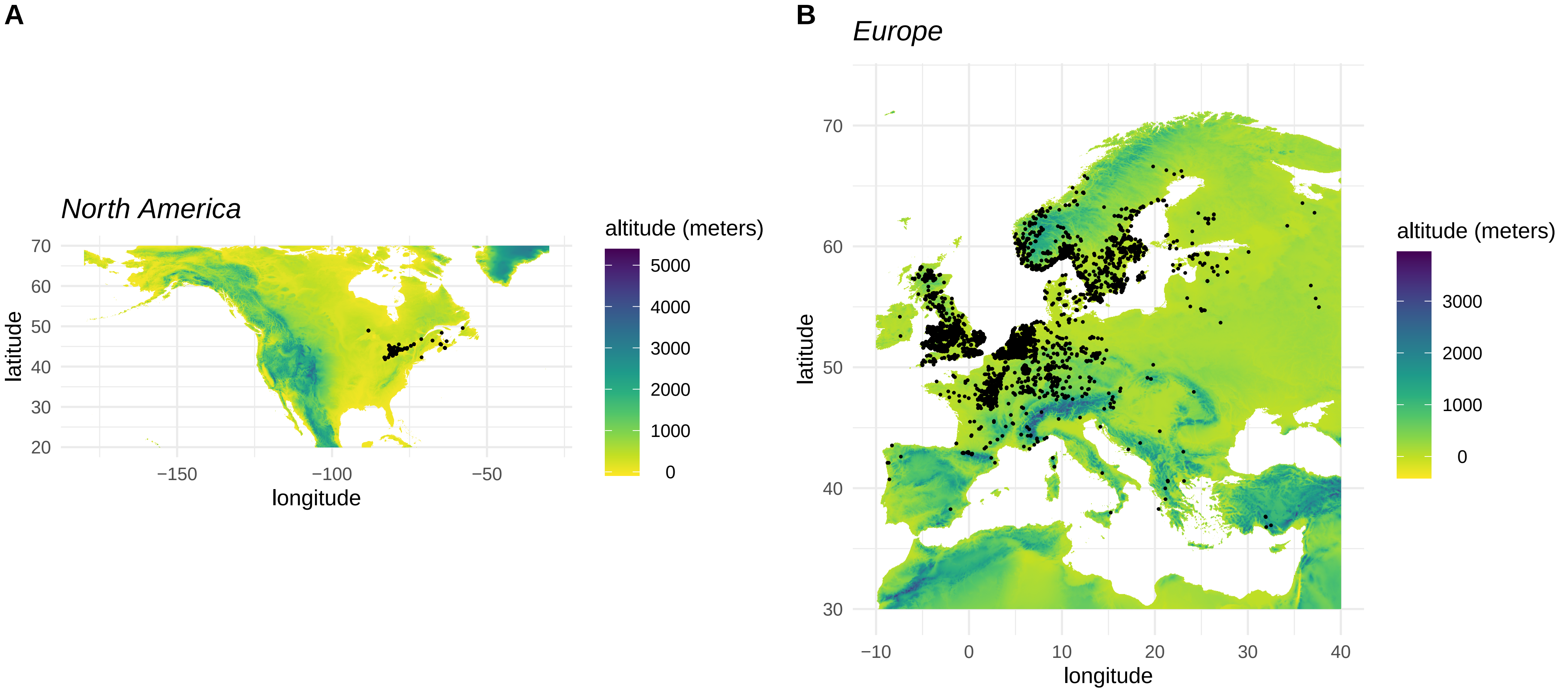

### Appendix S1c

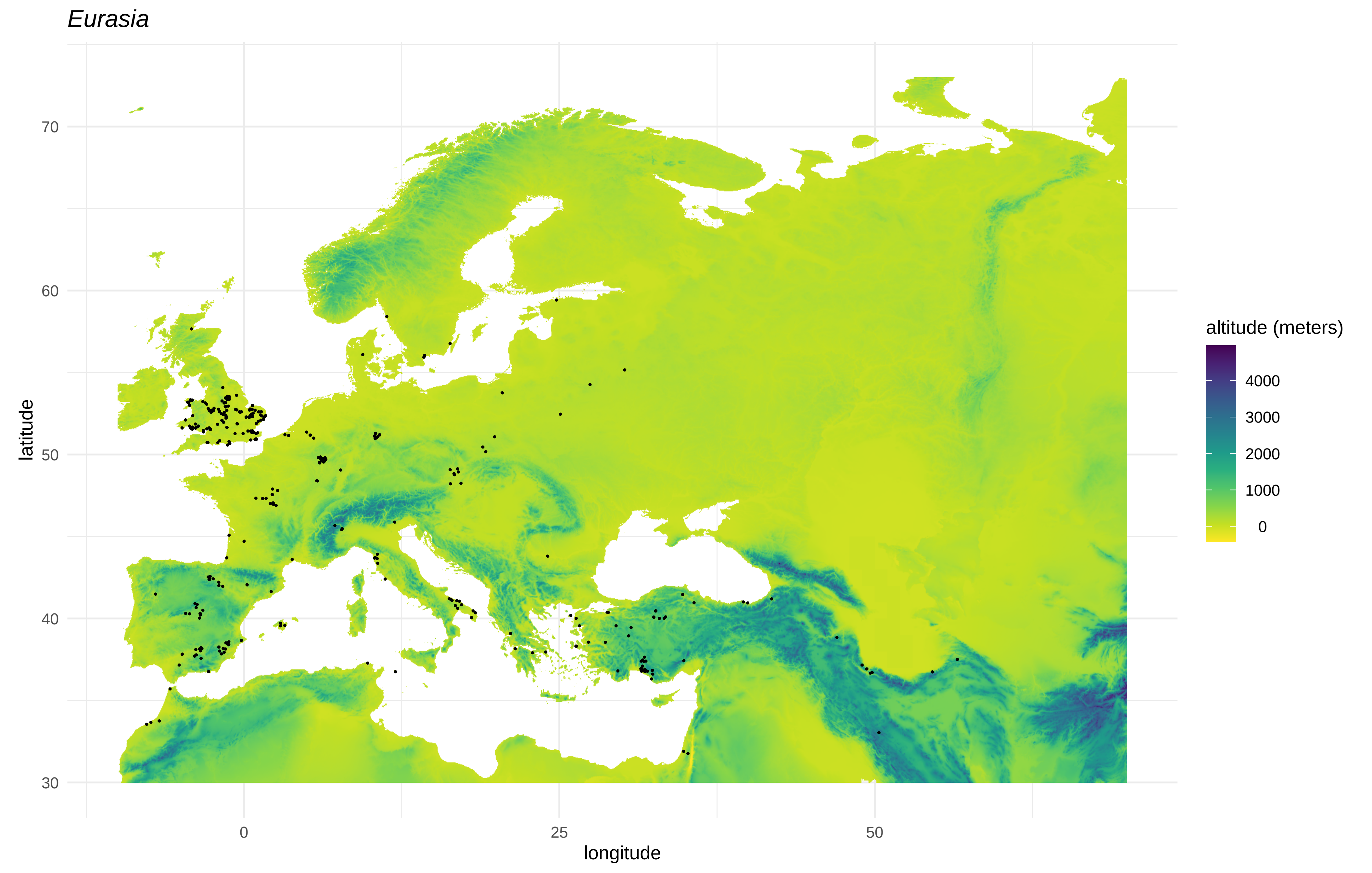

### Appendix S2

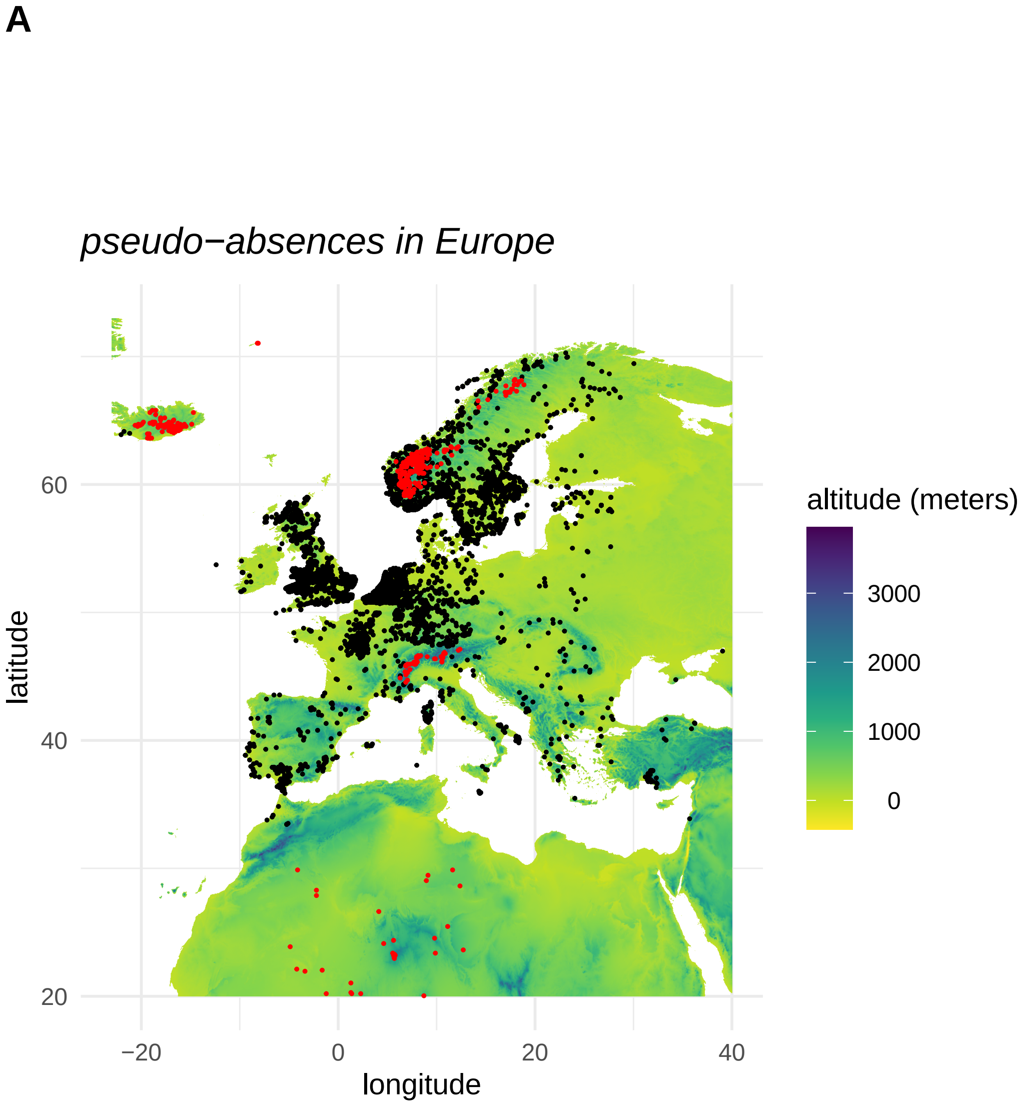

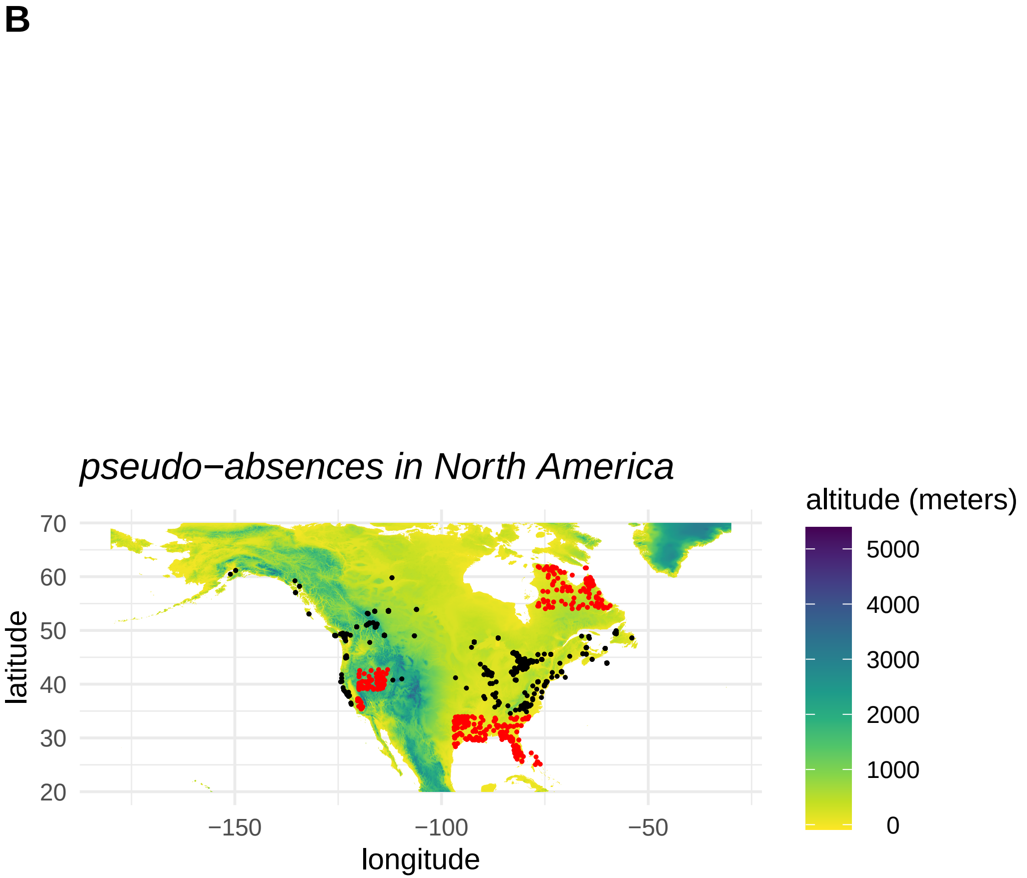

### Appendix S3

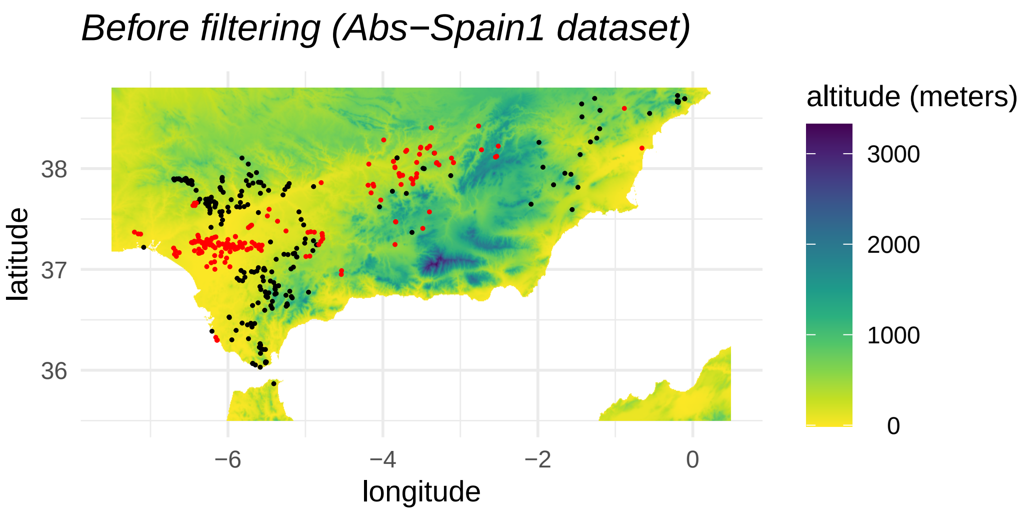

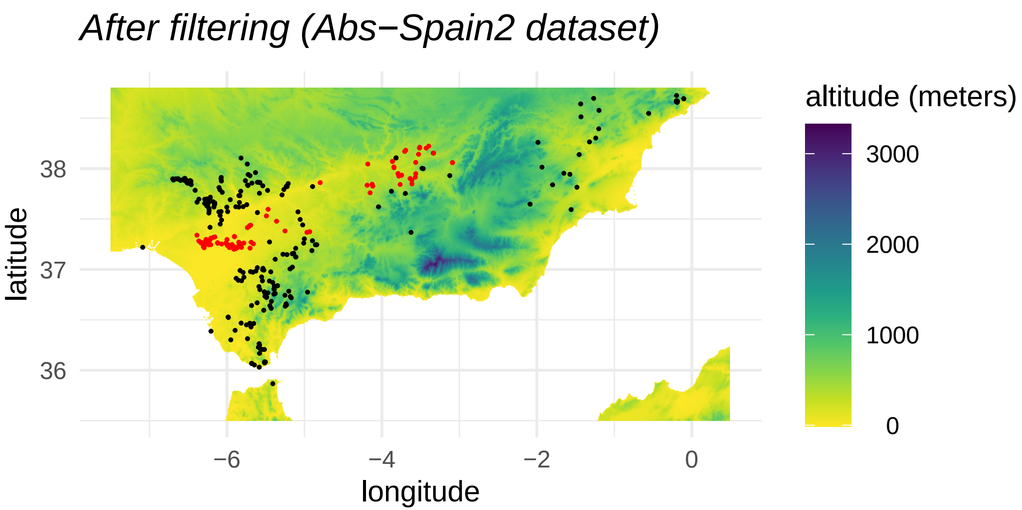

### Appendix S4

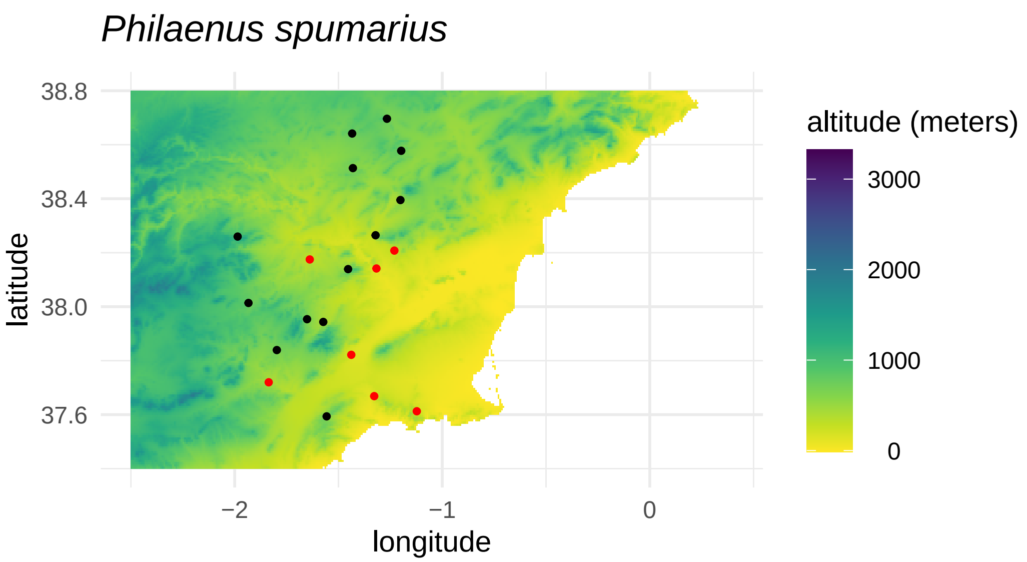

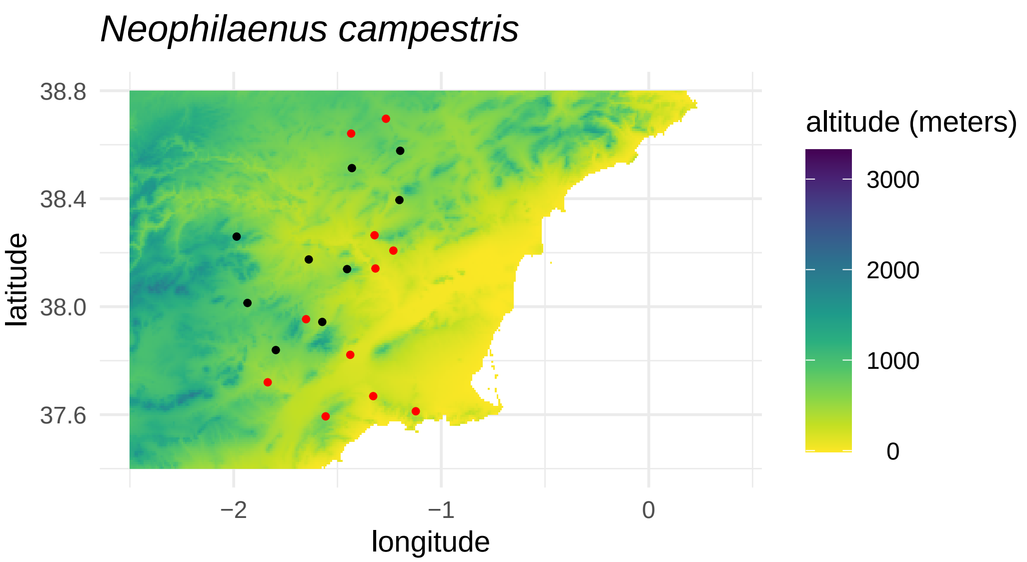
