## Appendix S1d for "The risk of *Xylella fastidiosa* outbreaks will decrease in the Mediterranean olive-producing regions"

**Appendix S1** Number of occurrences available to model the potential distribution of the main efficient or putative vectors of *Xylella fastidiosa* in Europe.

| **Species** | **Number of occurrence records before declusteing** | **Number of occurrence records after declustering** |
| --- | --- | --- |
| *Philaenus spumarius* | 17,618 | 535 |
| *Neophileanus campestris* | 473 | 139 |
| *Aphrophora alni* | 8,102 | 295 |
