## Appendix S5 for "The risk of *Xylella fastidiosa* outbreaks will decrease in the Mediterranean olive-producing regions"

**Appendix S4** Latitude and longitude range limits of the square selected for generating pseudo-absences when modelling the potential distribution of the main efficient or putative vectors of *Xylella fastidiosa* in Europe.

| **Species** | **Longitude range limits (in degrees)** | **Latitude range limits (in degrees)** |
| --- | --- | --- |
| *Neophileanus campestris* | -15; 60 | 30; 70 |
| *Aphrophora alni* | -100; 60 | 30; 70 |
